# Mutational Survivorship Bias: The case of PNKP

**DOI:** 10.1101/2020.08.03.233957

**Authors:** Luis Bermudez-Guzman, Gabriel Jimenez-Huezo, Andrés Arguedas, Alejandro Leal

## Abstract

The molecular function of a protein relies on its structure. Understanding how mutations alter structure and function in multi-domain proteins, is key to elucidate how a pathological phenotype is generated. However, one may fall into the logical bias of assessing protein damage only based on the mutations that are viable (survivorship bias), which can lead to partial conclusions. This is the case of PNKP, an important nuclear and mitochondrial DNA repair enzyme with kinase and phosphatase function. Most mutations in PNKP are confined to the kinase domain, leading to a pathological spectrum of three apparently distinct clinical entities. Since proteins and domains may have a different tolerance to disease causing mutations, we evaluated whether mutations in PNKP are under survivorship bias. Even when all mutations in the kinase domain are deleterious, we found a mayor mutation tolerability landscape in terms of survival. Instead, the phosphatase domain is less tolerant due to its low mutation rates, higher degree of sequence conservation, lower dN/dS ratios, and more disease-propensity hotspots. Thus, in multi-domain proteins, we propose the term “Wald’s domain” for those who are not apparently more associated with disease, but that are less resistant to mutations in terms of survival. Together, our results support previous experimental evidence that demonstrated that the phosphatase domain is functionally more necessary and relevant for DNA repair, especially in the context of the development of the central nervous system. Thus, this bias should be taken into account when analyzing the mutational landscape in protein structure, function, and finally in disease.

## Introduction

About two-thirds of the prokaryotic proteome and 80% of eukaryote proteins are multi-domain proteins [1]. It is possibly that proteomes have evolved from a limited repertoire of domain families, and multi-domain proteins were assembled from combinations of these [2]. Notably, proteins and domains differ in how well they tolerate mutations, suggesting that not all mutations are functionally important [3]. Disease-causing mutations frequently involve important changes in amino acid physicochemical properties that tend to destabilize proteins and their interactions [4]. These mutations occur mostly in the protein core, predominantly in helixes and coil regions, and not so frequently in beta strand structures [5]. Protein-protein interaction regions called interfaces, are also important “hotspots” for mutations. In fact, when compared to random segregation, disease-causing nonsynonymous single nucleotide polymorphisms are preferentially located at protein-protein interfaces rather than surface noninterface regions [6]. Within this interface, mutations are preferentially located in the “core” (residues solvent-inaccessible), as opposed to the “rim” (partially solvent-accessible). Interface rim is significantly enriched in polymorphisms, like the remaining non-interacting surface [7]. In addition, mutations in ligand-binding sites close to interfaces and residues related to enzymatic function are especially associated with disease [8].

Most mutations we observe in patients are those that manage to be viable despite affecting protein functioning. Mutations that result in unviable proteins and eventually unviable organisms, escape the analysis because we hardly see them in patients. In fact, it has been shown that proteins and domains vary in their tolerance to non-synonymous point mutations according to their mutation rates, sequence conservation and interaction networks [3]. This scenario resembles the one found by the mathematician Abraham Wald during World War II. At that moment, the Center for Naval Analyses (CNA) wanted to determine which sites should be reinforced on their planes based on the damage the planes had once they returned from combat. Wald pointed out that CNA was only considering the planes that had survived their missions for this purpose [9]. Since the planes that had been shot down were not present for the damage assessment, the holes in the planes that returned represented areas where an airplane could be damaged and still safely return (Fig. 1). Wald proposed that the Navy reinforce the areas where the returning planes were undamaged since those were the areas that, if attacked, would cause the plane to be lost.

**Figure 1.**
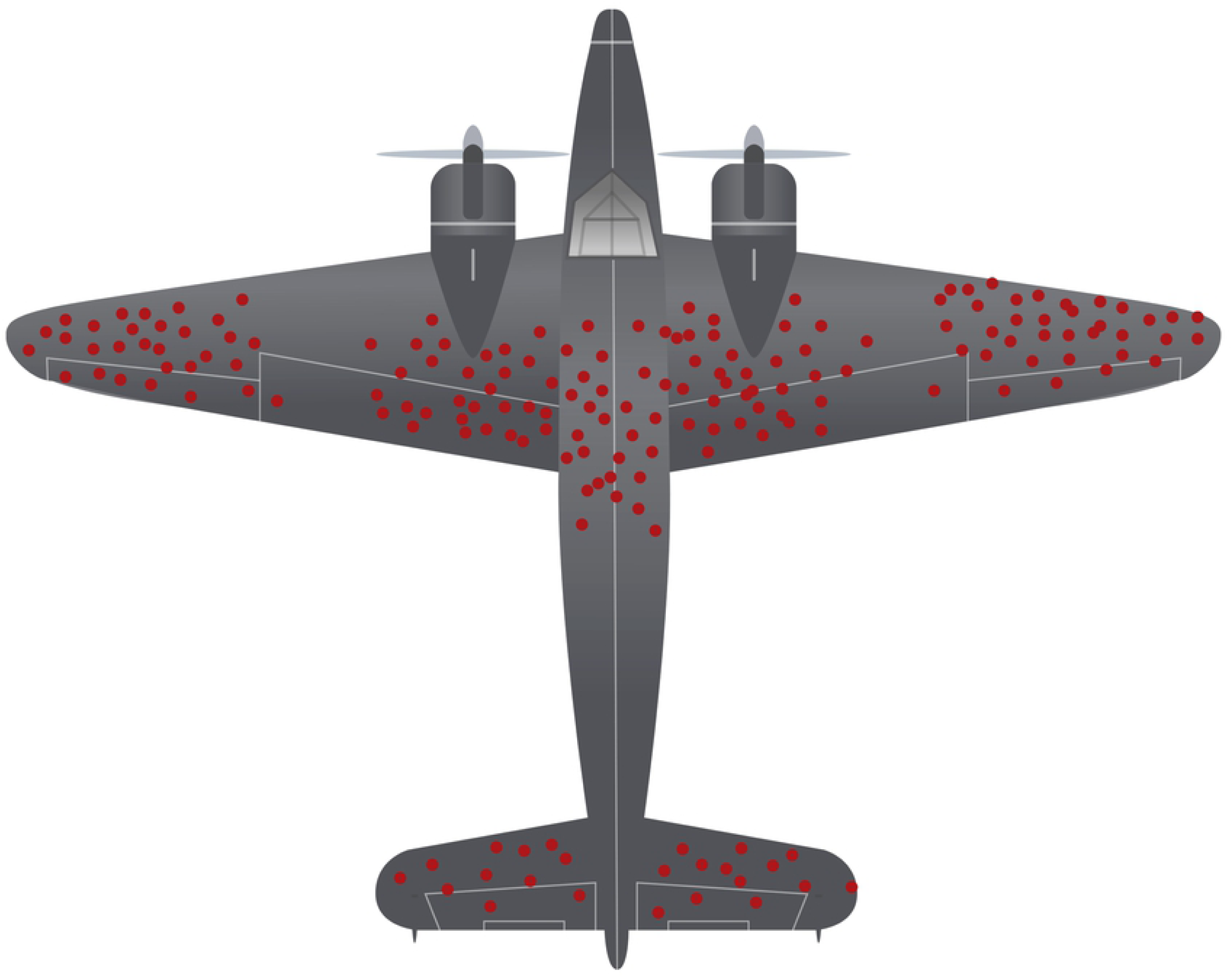
Depiction of Wald’s idea regarding survivorship bias and the overlooking of those events that did not succeed in a selection process. The red dots represent hypothetical areas where an airplane could be damaged and still return.

Another interesting case was the one that occurred during World War I. After the introduction of the Brodie helmet and others [10] it is said that there was an increase in hospitalizations for head injuries. Is not that wearing a helmet caused more injuries, but fewer deaths. Soldiers who previously died for head trauma were now surviving so they could be taken to a field hospital. This observation was later called survivorship bias, as the logical error of focusing on the events that made it past some selection process and overlooking those that did not. This concept is not new since different types of biases have been proposed historically in genetics and evolution. One example of this is the so-called transition bias, as the pattern in which nucleotide transitions are favored several times over transversions, attributed to the fact that transitions are more conservative in their effects on proteins [11].

Analyzing the pathogenicity of the mutations according to the different domains of a protein can provide significant information about the sites where it is more likely to find disease-causing mutation [12]. Thus, based on Wald’s observation, we believe that many proteins could be subject to survivorship bias. In other words, we believe that in genotype-phenotype correlation, we are only considering disease-causing mutations that are tolerated when assessing protein damage. One of these proteins could be the Polynucleotide Kinase 3’-Phosphatase (PNKP), an important nuclear and mitochondrial DNA repair enzyme [13]. This protein consists of 521 amino acids with three characterized domains: an N-terminal FHA (ForkHead-Associated) domain, the phosphatase, and the C-terminal kinase [14]. PNKP is recruited to the damage site through the interaction between its FHA domain and the X-Ray Repair Cross Complementing one protein (XRCC1), for Single-Strand Break Repair (SSBR) [15]. It can be also recruited by XRCC4 for Double Strand Breaks (DSBs) as part of the Non-Homologous End Joining pathway [16].

As the molecular function of a protein depends on its three-dimensional structure, determining the protein structure constitutes an important approach to understand the effect of mutations at the functional level [17]. When a mutation that causes single amino acid variation (SAV) is discovered, the general pipeline is to analyze its pathogenicity in terms of amino acid conservation, protein structural features and even gene ontology [18]. However, the degree of conservation of the residue is not always informative since it has been shown that disease-causing missense variants also affect very variable sites [19]. Usually, if there are no previous experimental studies, we try to analyze different aspects *in vitro* such as the amount of protein, mutant protein stability, and function. However, in many cases, it is very difficult to analyze all the mutations and their effects on protein structure and to what extent they can generate one or more pathological phenotypes.

Mutations in PNKP are associated with a wide pathological spectrum. The most severe phenotype is Microcephaly with early-onset Seizures (MCSZ) [20]. In the middle, Ataxia with Oculomotor Apraxia 4 (AOA4) [21] and in the other end the milder Charcot-Marie-Tooth disease type 2B2 (CMT2B2) [22–24]. The vast majority of mutations in PNKP are confined to the kinase domain, both in homozygous and compound heterozygous conditions. To date, it is not understood how mutations in a single domain can cause such a broad clinical spectrum, so it is speculated that the kinase domain is the one that determines the degree of pathogenicity. In this paper, we analyze the mutations observed in each domain of PNKP in order to determine whether they are subject to mutational survivorship bias or not. To do this, we combine *in silico* analysis, published experimental data and open sequencing data to combine both structural and functional information.

## Results

### Multiple sequence alignment

First, we wanted to analyze some protein features of PNKP. From all the available protein sequences in the Uniprot database, seventy sequences of twenty taxonomic groups were selected. We performed a Multiple Alignment using Fast Fourier Transform (MAFFT) against the human PNKP sequence (see Methods for the ID number of each sequence). We generated a phylogenetic tree using this alignment with the ITOL platform [25]. We also ran a BlastP analysis to determine the percent identity among species and we noticed that human PNKP has low identity compared to some higher groups (Fig. 2).

**Figure 2.**
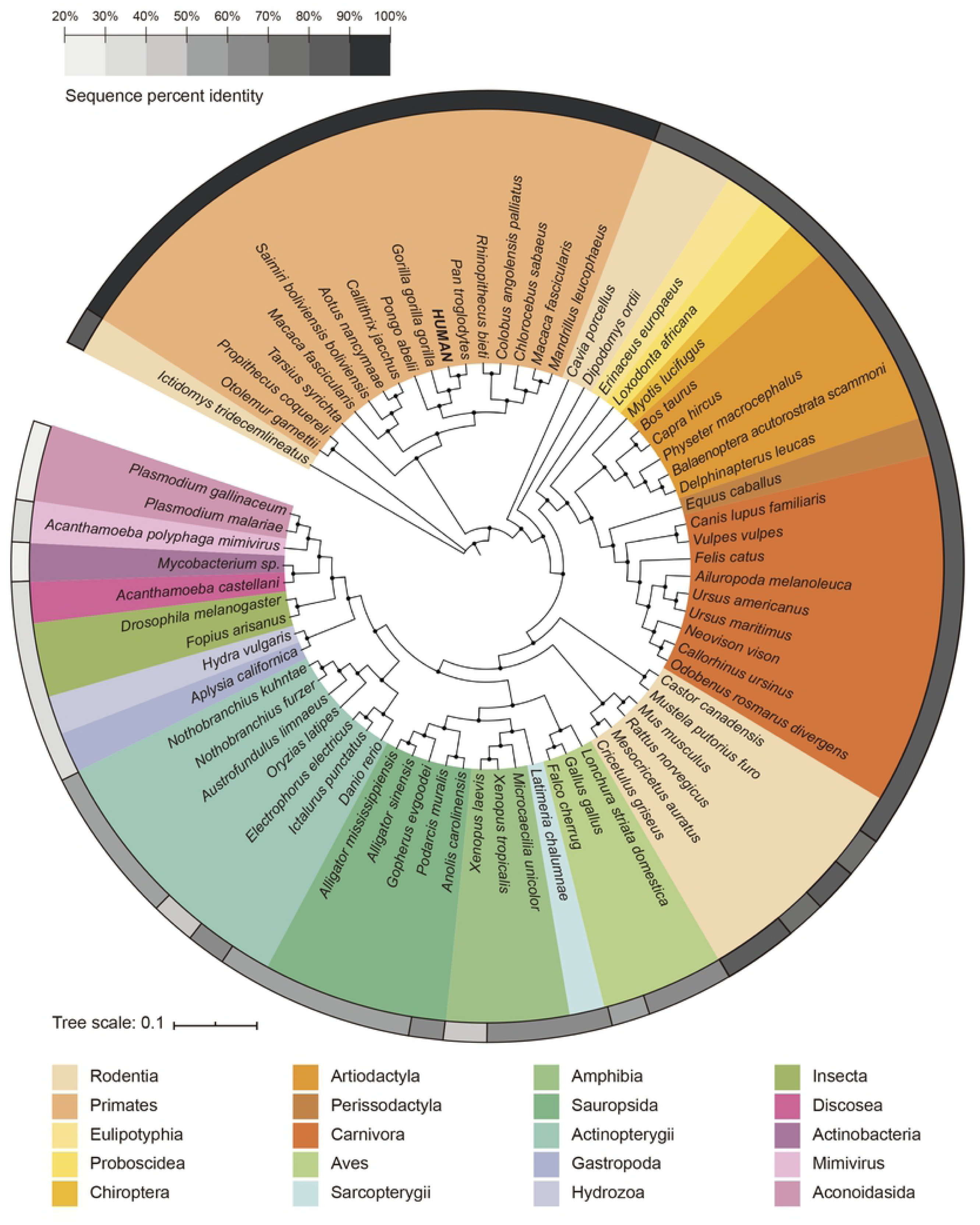
Phylogenetic tree of the seventy species used in the multiple sequence alignment according to their taxonomic group. The gray circular scale represents the percent identity with respect to the human protein sequence of PNKP.

As expected, the group of primates was the closest to the human, with identities between 90-100%. The species belonging to the other eight orders of mammals presented a range between 80-90% (e.g. *Mus musculus*: 80.88%). The species of the group of birds, reptiles, and amphibians had a range between 50-70% (e.g. *Gallus gallus*: 66.19%). The fish group showed a lower range 50-60%. The rest of the species had percent identities below 50% (*e*.*g*. Drosophila melanogaster: 40.16%).

### Percent identity analysis of SSBR proteins

To further confirm the low percent identity of the human sequence of PNKP, we compared the degree of identity between the sequences of different enzymes from the Single-Strand Break Repair (SSBR) pathway with respect to ten vertebrate species. Human protein sequences of PNKP, PARP1, XRCC1, POLB, and LIG3 were analyzed using the BlastP tool [26]. We identify the percent identity of *Pan troglodytes, Equus caballus, Sus scrofa, Canis lupus, Felis catus, Mus musculus, Rattus norvegicus, Gallus gallus, Xenopus laevis*, and *Danio rerio* with respect to the corresponding human sequence (see Methods for the ID number of each sequence).

We later used the percentage corresponding to the difference in the sequence identity between each species and the human sequence. The protein sequence that showed the lowest difference between species was POLB, followed by PARP1, LIGIII, XRCC1 and finally PNKP with the lowest similarity among species (Fig 3). Surprisingly, the difference between the human PNKP sequence and *Mus musculus* (mouse) was almost 20%. To further evaluate this difference, an ANOVA test was performed and we confirmed that there is a smaller gradient, and thus less differences between species, for POLB protein sequence, and a larger gradient, with a notable difference in sequence percent identity between species, for PNKP and XRCC1 (F value = 13.368; p-value = 8.9 ×10-7) (Fig. 4).

**Figure 3.**
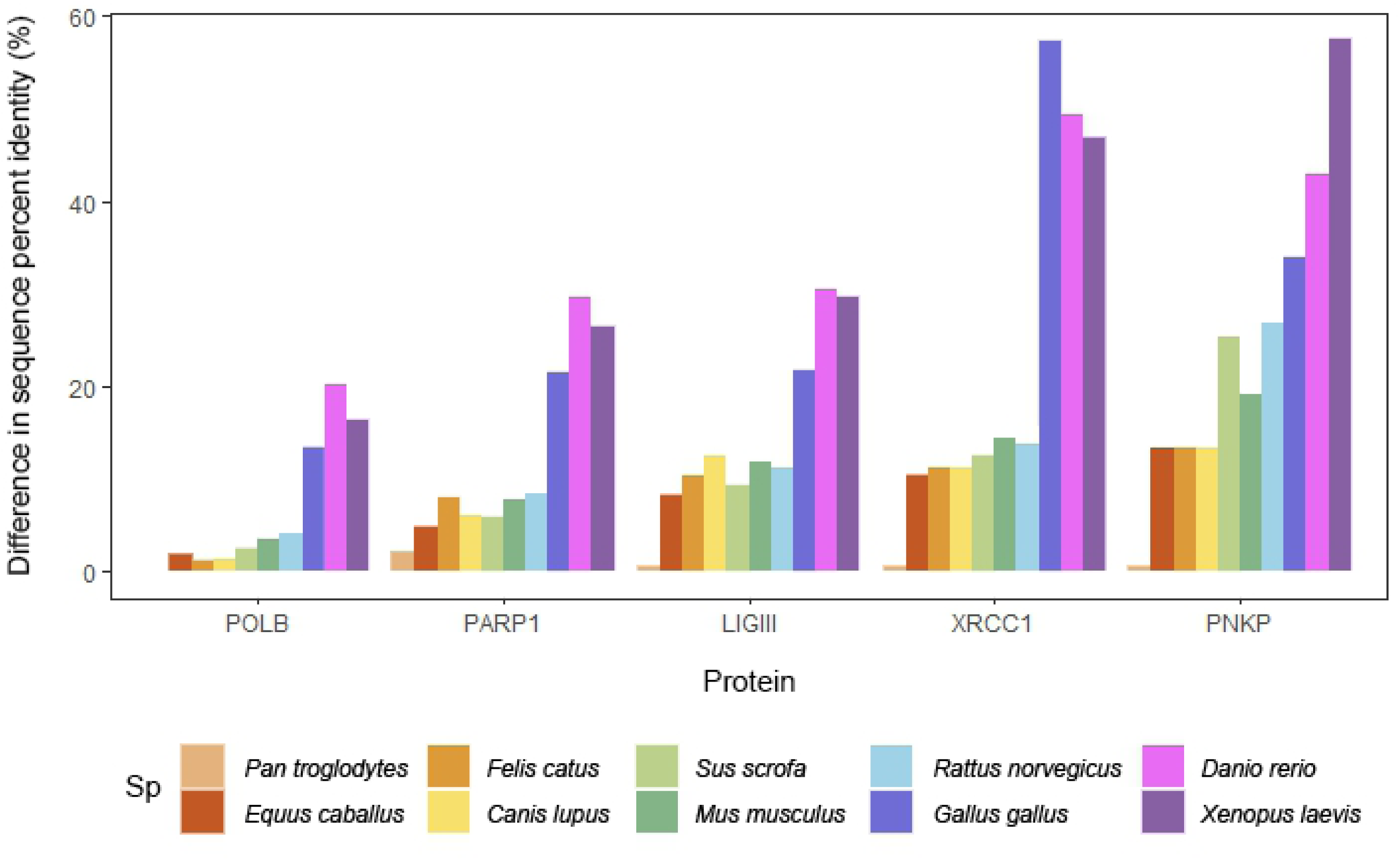
Comparison of the difference in the percent identity between the human sequence of each SSBR protein and several species. The higher the bar, the more different is the sequence of each animal with respect to the human.

**Figure 4.**
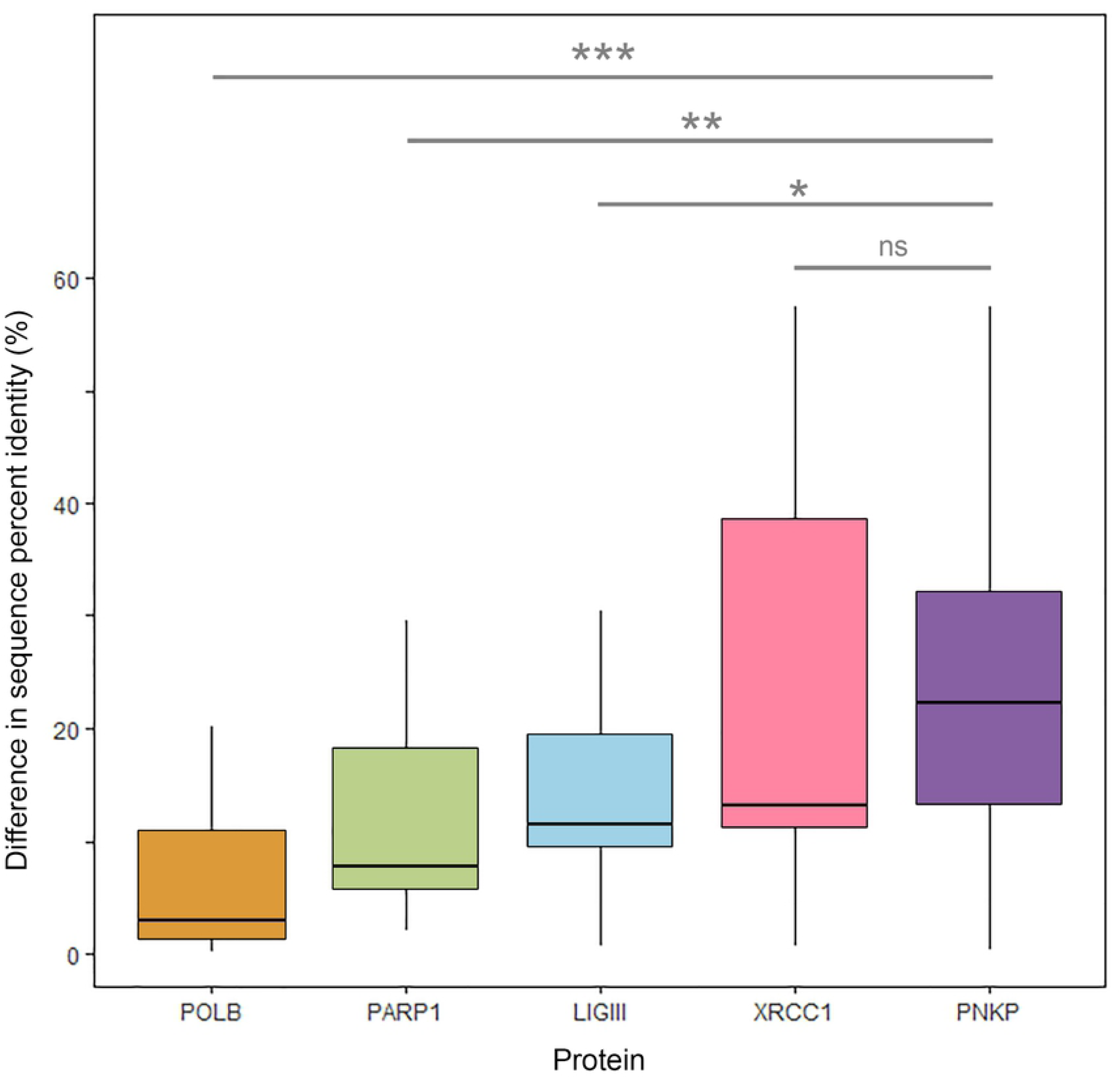
Difference in the percent identity of each SSBR protein between species compared to the human sequence (* p<0.05; ** p<0,01; *** p<0,001; ns: not significant).

Using Tukey’s intervals for multiple comparisons, we observed that POLB presented lower differences among species than PNKP and XRCC1 (p < 0.05), while no differences were found with respect to PARP1 and LIGIII (p > 0.05). No significant differences were found between PARP1, LIGIII and XRCC1 (p > 0.05). PNKP shows bigger differences in sequence identity than POLB (p =0.0000035), PARP1 (p = 0,001) and LIG III (p = 0.013). No difference was found between PNKP and XRCC1 (p = 0,97). These differences could be related to the evolutionary origin and mutational rates of each domain of PNKP. It seems that the phosphatase domain is more conserved among different species, while the kinase presents homology with different enzymes (see Supplementary Figure 1).

### Conservation analysis

As we noticed the difference between the evolutionary history of each domain, we wondered if the degree of sequence conservation was higher in the phosphatase than in the kinase domain. The previous MAFFT alignment was used as input in the ConSurf 2016 platform to get and compare the conservation scores of the amino acids among the phosphatase (from amino acid 146 to 337) and kinase domain (from 341 to 521) (Fig. 5a). There was a total of 192 observations for the phosphatase domain and 181 for the kinase, corresponding to each amino acid. This makes the results using absolute numbers almost equal to those using relative numbers. Homogeneity chi-squared test showed that both domains have different conservation scores (□^2^ = 22.865, df = 8, p = 0.0035, Fig. 5b), with higher conservation in the phosphatase domain. Specifically, in the cases of conservation scores 8 and 9, the percentage of observations from phosphatase domain was 34.4%, while, for kinase this percentage was only 16.99%. On the other hand, for lower conservation scores from 1 to 3, the percentage in phosphatase was 14.59%, while for kinase was 24.94%. ConSurf also predicted the 3D structure of PNKP by using Modeller and HHPred Model algorithms to color the structure according to the conservation scores (Fig. 5c). Interestingly, the region that joins the FHA domain with the phosphatase, has a very low conservation score.

**Figure 5.**
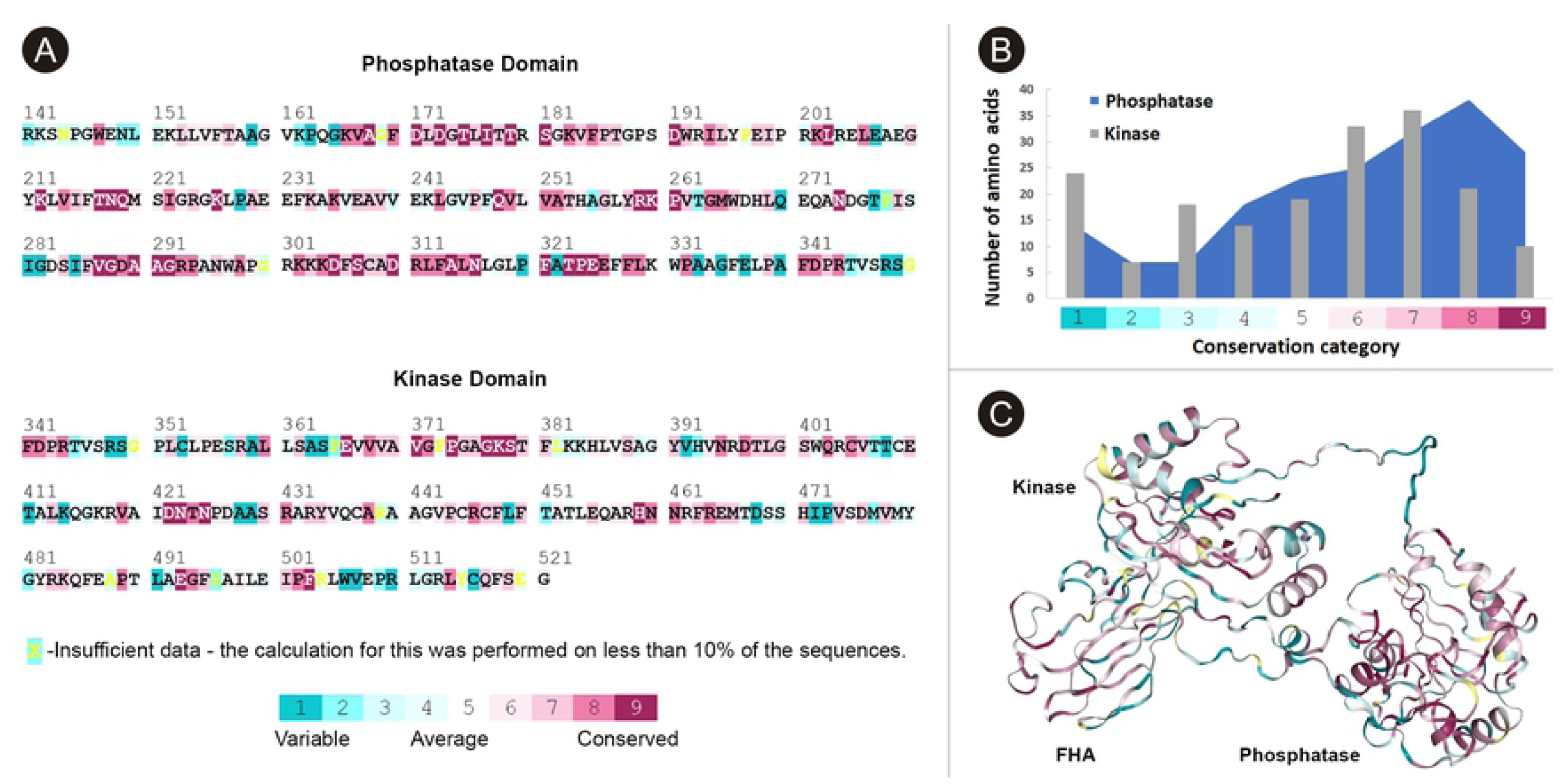
Conservation analysis of the phosphatase and kinase domains. A) Color conservation score for the phosphatase and kinase domain sequence according to ConSurf 2016. B) Number of amino acids according to the conservation score for each domain. C) Prediction of the colored 3D structure according to the degree of conservation between each domain. The random coil linking the different domains show a low conservation score, while the degree of greater conservation is in the phosphatase domain.

### PNKP 3D structure prediction

Next, to analyze the structural damage of mutations in PNKP, several platforms were used to predict its 3D structure since there is no structure of human PNKP in the RCSB Protein Data Bank. Swiss-Model generated the best model using only two templates (3zvn.1.A, 2brf.1.A), however, it was not able to generate the complete structure as it did not find a template to model the sequence between the FHA domain and the phosphatase, generating a gap in the 3D model. Interestingly, this is the same region with a very low conservation score from Figure 5c. For this reason, this model was not included in the evaluation. Since ConSurf generated its own model using the Modeller platform, we included this model to be evaluated alongside the others. Modeller used six different templates (3zvl_A, 1ujx_A, 1yj5_C, 2brf_A, 3kt9_A, 1ly1_A) and Phyre2 selected 7 templates (d2brfa1, d1ujxa_, c1yj5B_, c3zvmA_, d1yjma1, d1qu5a_, c1yj5C_) while I-TASSER used ten templates to model the structure (1yj5B, 1ujxA, 3zvlA, 2brfA, 1yj5B, 1yj5, 1yjmA, 1yj5, 1yj5B, 3kt9A). From all the models generated with I-TASSER, we chose the model with the lowest C-score (−2.22).

The Ramachandran plots of each model were generated with the Swiss-Model Structure Assessment tool (Fig. 6). The core regions (dark green in the figure) contain the most favorable combinations of φ and ψ (torsional angles). The allowed regions (light green in the figure) can usually contain fewer data points than the core regions. The generous regions (remaining colored area in the figure) extend beyond the allowed regions. Dots within white spaces are generally considered outliers. According to Ramachandran plots, the model generated by Modeller presented the highest number of amino acids in a favored position and the lowest number of outliers (Fig. 6A). The comparison of each model with a non-redundant set of PDB structures showed that this model has the lowest value for the Qualitative Model Energy ANalysis (QMEAN) (−4.34) (Fig. 6B). This indicates whether the QMEAN score of the model is comparable to what one would expect from experimental structures of similar size. The QMEAN for the model generated in Phyre2 was −7.06 and for I-TASSER was −10.21 (QMEAN Z-scores around zero indicate good agreement between the model structure and experimental structures of similar size). The “Local Quality” plot shows for each residue of the model (reported on the x-axis), the expected similarity to the native structure (y-axis) (Fig. 6C). Typically, residues showing a score below 0.6 are expected to be of low quality. In this case, Modeller presented the highest number of values above this threshold. More detailed results of all the rubrics evaluated in MolProbity for each model are available in Supplementary information (Supplementary Table 1). To compare the overall quality of this result, we also modelled the structure of PARP1, since it has more crystal structures reported in RCSB PDB. The MolProbity Score of this model was 3.3, similar to the one we obtained with our model (3.8) (data not shown). In addition, we performed a structural alignment between our 3D structure generated with Modeller and the mouse Phosphatase-Kinase crystal structure of PNKP (PDB ID: 3ZVL). Notably, both structures were almost equal (TM-score: 0.99357; RMSD: 0.25 Å).

**Table 1.** Structural analysis of missense mutations reported in the kinase (orange) and phosphatase (purple) domain of PNKP.

| **Mutation** | **Conservation Score (1-9)** | **SuSPect (0-100)** | **Hotspot** | **Binding Site** | **$\Delta\Delta G$ (Kcal/mol)** | **Structural Damage** |
| --- | --- | --- | --- | --- | --- | --- |
| p.Arg462Pro | 6 | 23 | No | Nearby | 0.201 | Cavity expansion |
| p.Lys378Thr | 9 | 92 | Yes | Yes | -0.957 | Cavity expansion |
| p.Gly375Trp | 6 | 86 | Yes | Yes | 1.270 | No |
| p.Pro343Leu | 7 | 36 | No | No | 1.543 | No |
| p.Gly442Ser | 5 | 9 | No | No | -1.822 | No |
| p.Leu399Pro | 7 | 64 | No | No | -1.661 | Secondary Structure |
| p.Cys409Trp | 3 | 71 | No | No | 0.059 | H-Bond breakage |
| p.Leu176Phe | 7 | 87 | Yes | Yes | 0.188 | No |
| p.Gly292Arg | 8 | 37 | No | No | 1.063 | Buried Gly replacement |
| p.Glu326Lys | 8 | 28 | No | No | -0.788 | No |

**Figure 6.**
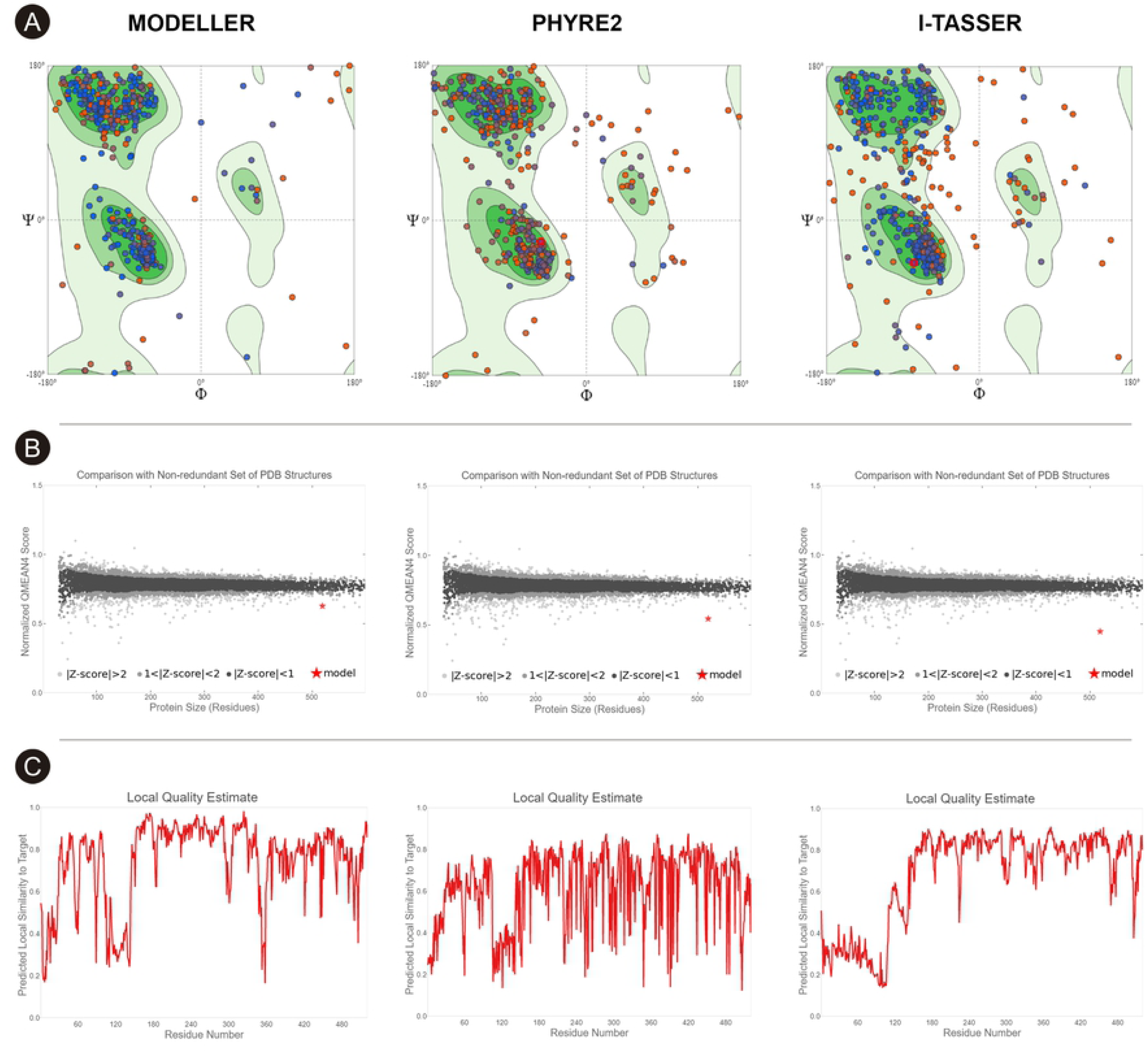
Structure assessment of the PNKP 3D models. A) Ramachandran plots of the three models generated on each platform. B) Comparison of the model with a non-redundant set of PDB structures. C) Local quality estimate according to the local similarity to the target.

### Prediction of pathogenicity of mutations in PNKP

Once we determined that the phosphatase domain is more conserved and generated the best 3D structure of PNKP, we used the SuSPect platform [18] to analyze the mutational sensitivity of the amino acids belonging to each domain. As input, we used the structure generated in Modeller and the FASTA sequence, as this method predict how likely single amino acid variants (SAVs) are to be associated with disease. First, the average of the 20 obtained scores for each amino acid was calculated, and the distribution of said averages was plotted. As we noticed that values presented an asymmetric distribution, with most positions having an average disease susceptibility lower than 50, we plot the average mutation susceptibility of each amino acid. As a threshold, we determined the critical value as the 95^th^ percentile, close to a disease-propensity score of 75 (Fig. 7). This means that, for a position to be considered as part of a “disease-propensity hotspot”, the region must present an average higher than 75 for all possible single amino acid variants (Fig. 8).

**Figure 7.**
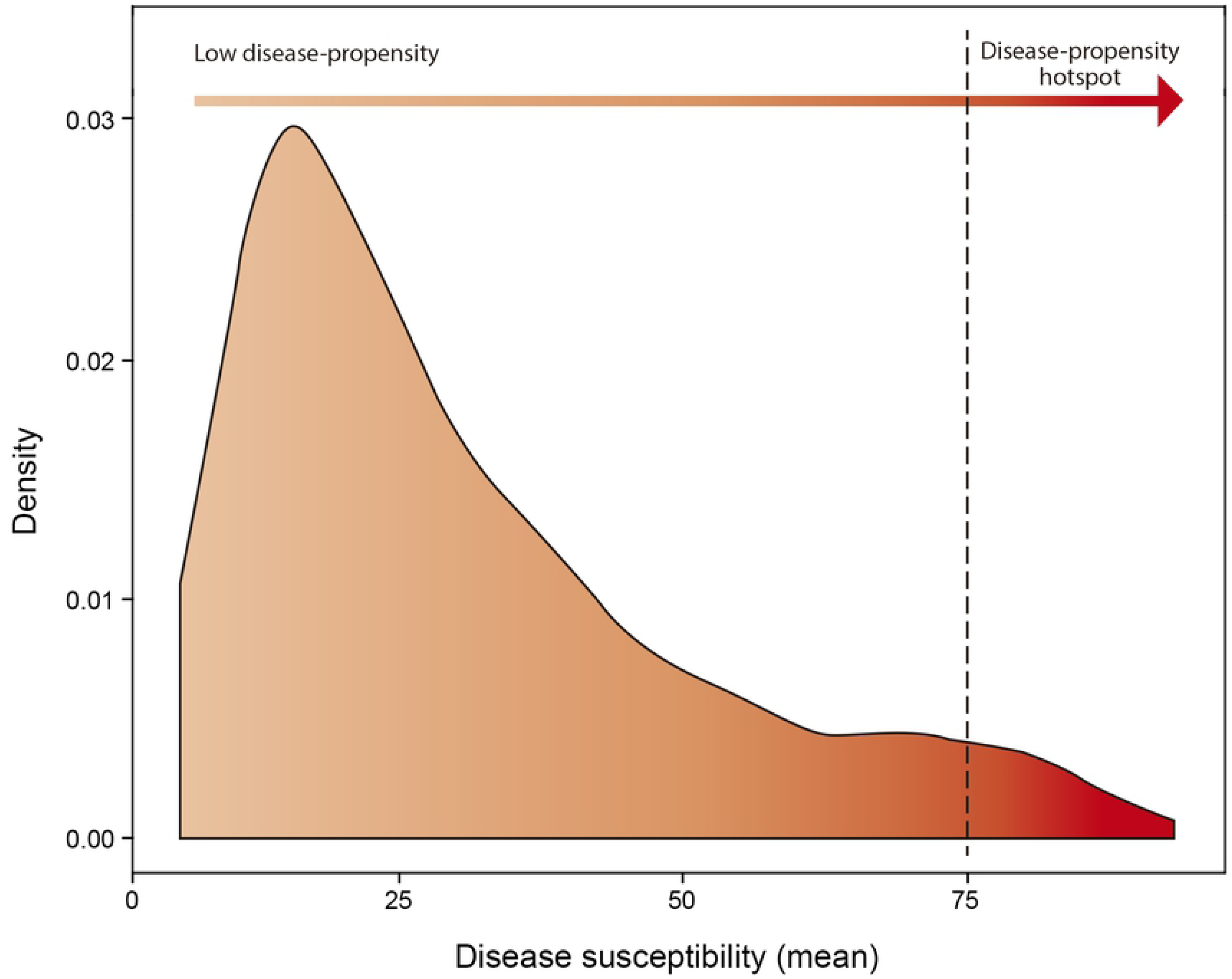
Distribution of the average values for disease susceptibility (SuSPect) scores for all the amino acids of PNKP.

**Figure 8.**
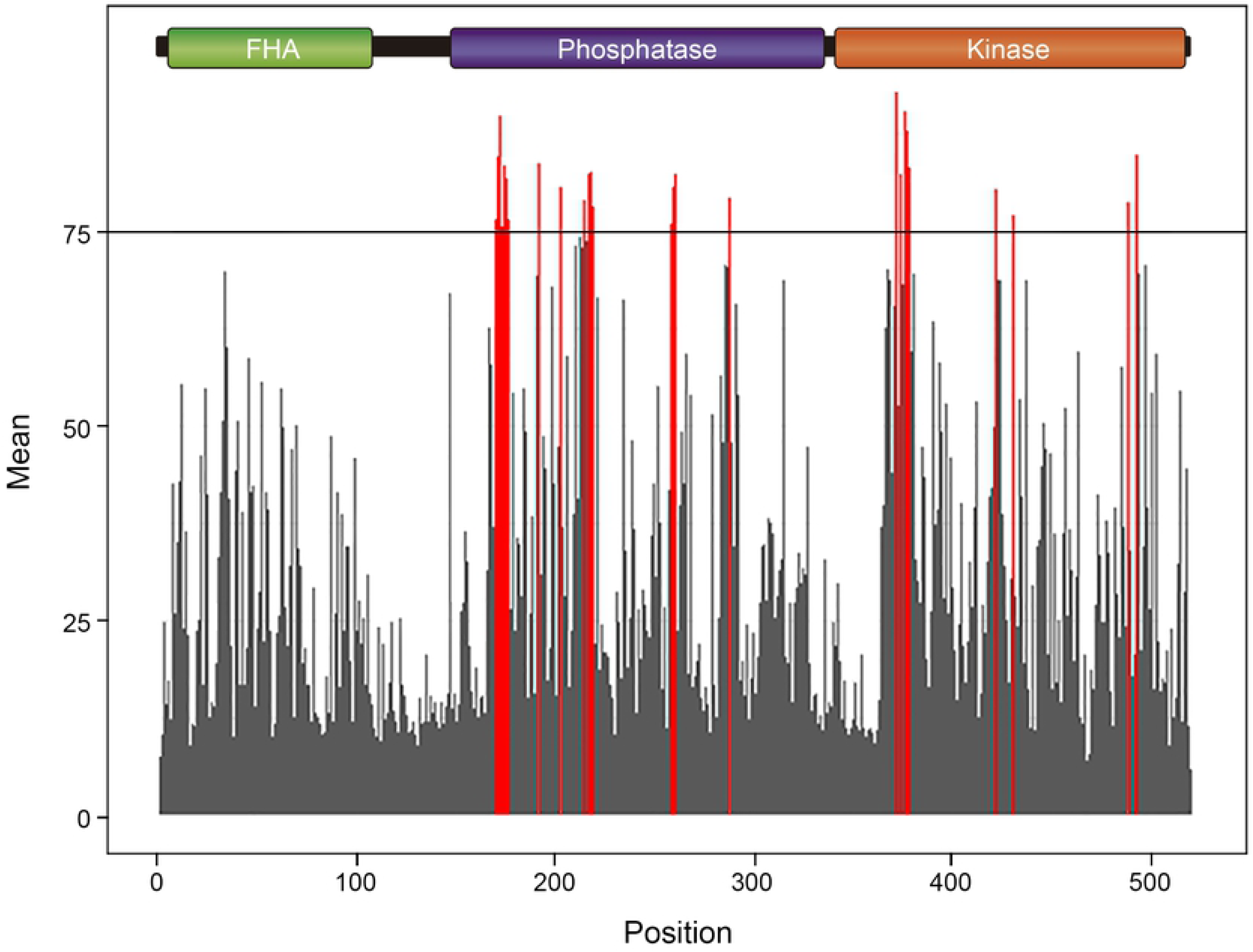
Analysis of disease-propensity hotspots. The average of disease susceptibility scores for each amino acid of PNKP was plotted. All variants with an average beyond the critical value of 75 are colored in red, but only consecutive positions with averages over the critical value are considered disease-propensity hotspots.

The only consecutive positions with averages over the critical value were from 170 to 177, from 217 to 219, and from 259 to 261 in the phosphatase domain. In the kinase domain only amino acids from 377 to 379 exceeded the threshold. Since this is an average, individual values could be much higher for some of the said positions, but this clearly indicated the high propensity to disease.

### The PNKP ligand-binding sites prediction

To confirm if disease-propensity hotspots correspond to functional ligand-binding sites in PNKP, we used COACH [27] and COFACTOR [28]. For each predicted site, the residues with the highest C-score (confidence score of the prediction from 0 to 1, where a higher score indicates a more reliable prediction) were chosen. For the DNA-binding site, the highest C-Score was given by the COFACTOR (0.61) for residues D171, N218, Q219, M220, R259, F306 (Fig. 9B) while COACH had a slightly lower C-Score (0.57) for the same residues but also including V184, F185, S221, R224, K226, A255, M477. Some of these residues coincide with the hotspots we found from amino acids 170-177 and 259-261 above (Figure 8). Only COACH predicted the ATP/ADP-binding site in the kinase domain, with a C-Score of 0.15 and the following residues: G375, A376, G377, K378, S379, T380, N460, F463, R464, F503, R504, L505, W506, Y515 (Fig. 9A). This site also coincides with the kinase disease-propensity hotspot from 377-379. For the phosphate group-binding site (the phosphate group of DNA), COFACTOR predicted with greater C-Score (0.42) the residues L172, D173, T217, N218, K260 (Fig. 9C). This site also corresponds to disease-propensity hotspots we found in the phosphatase domain from amino acids 170-177 and 259-261. COACH also predicted three amino acids for the Mg+2 binding sites (D171, D173, D289) which also matched a hotspot within the phosphatase domain (C-Score: 0.20, not show in the figure). We could not predict the kinase DNA-binding site. However, it was demonstrated that the DNA backbone approaches the kinase domain near a positively charged surface including R403, R482 and R483 residues [29]. Interestingly, these sites presented low average SuSPect values (R403 = 14.85; R482 = 39.35; R483 = 25.8).

**Figure 9.**
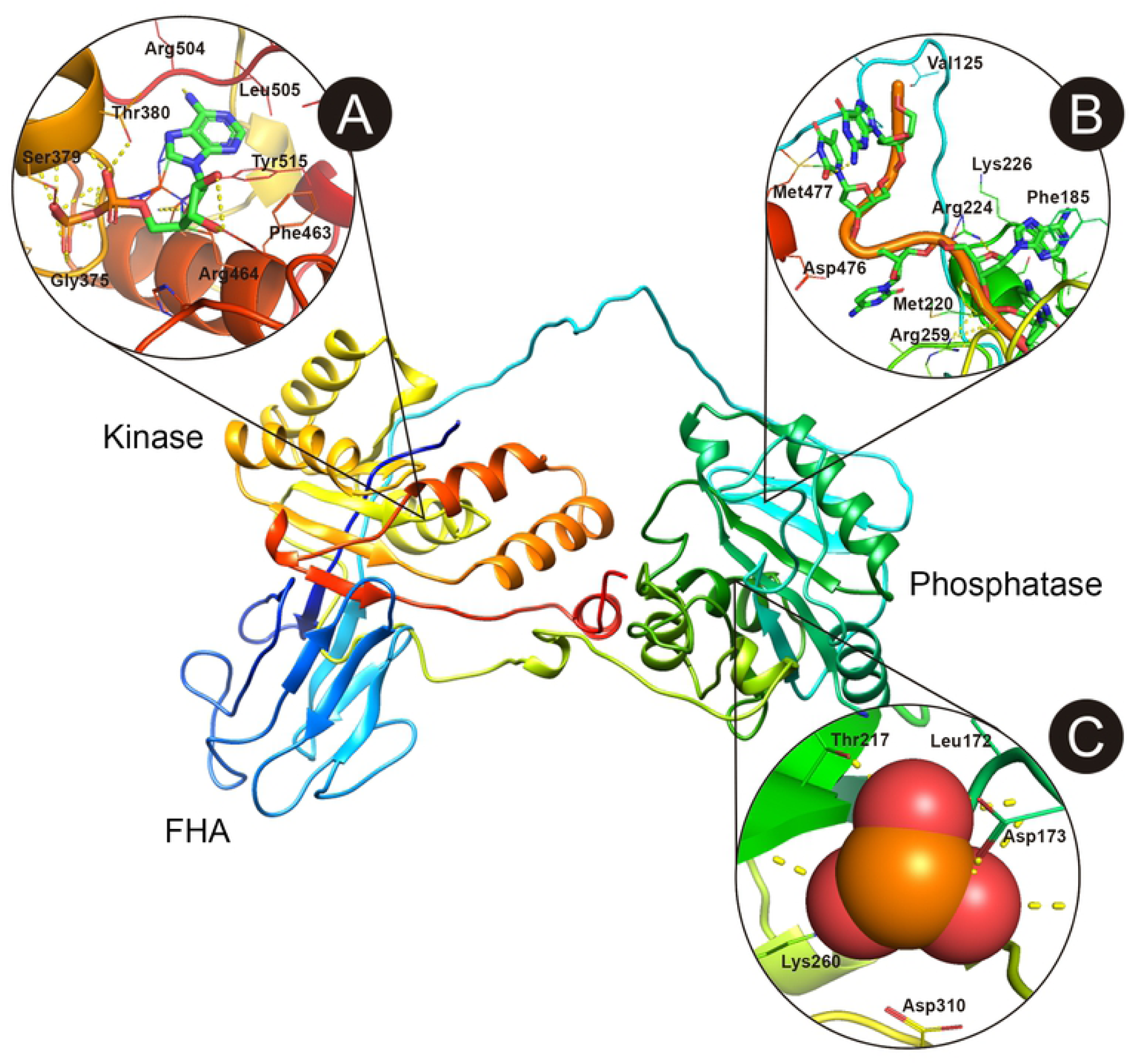
3D structure of PNKP (Modeller) showing the predicted most important ligand-binding sites. A) The binding site for ATP/ADP in the kinase domain. B) DNA binding site in the phosphatase domain. C) Phosphate group binding site in the phosphatase domain. Figure was elaborated with PyMol 2.3.

To add one more layer of complexity, we analyzed the potential post-translational modification (PTM) sites of PNKP. NetPhos 3.1 predicted S47, S114, T118, S126, S143, S221, S280, S307, T323, S347, T407 as phosphorylation sites (prediction score > 0.90). To confirm this prediction, we reviewed the Phospho.ELM and PhosphoSitePlus v6.5.9.1 database, which contain *in vivo* and *in vitro* phosphorylation data. Phospho.ELM confirmed the phosphorylation of S114, T118, T122, S126. Instead, PhosphoSitePlus also reported S12, T109, T111, S114, T118, T122, S126, N144, S143, G188, S364 and S379, although some with few references. Other PTMs were reported, as acetylation in K226, K233 and K242. Ubiquitination sites were also reported in K142, E151, K183, K226, and K484. However, these sites do not coincide with disease-propensity hotspots nor mutation sites.

### Structural damage of PNKP mutations

As we found that the disease-propensity hotspots matched some predicted ligand binding sites, we evaluated whether Single Amino acid Variants reported in patients with PNKP mutations also coincided with these hotspots. We also evaluated the structural damage of each mutation in the Missesense 3D platform using the structure predicted with Modeller. Dynamut were also used to analyze the impact of mutations on protein dynamics and stability resulting from vibrational entropy changes. We included the following variants: Arg462Pro, Lys378Thr, Gly375Trp, Leu399Pro, Cys409Trp, Pro343Leu, Gly442Ser, Leu176Phe, Gly292Arg and Glu326Lys (Fig. 10). Notably, the majority of these residues are located within the core of the protein.

**Figure 10.**
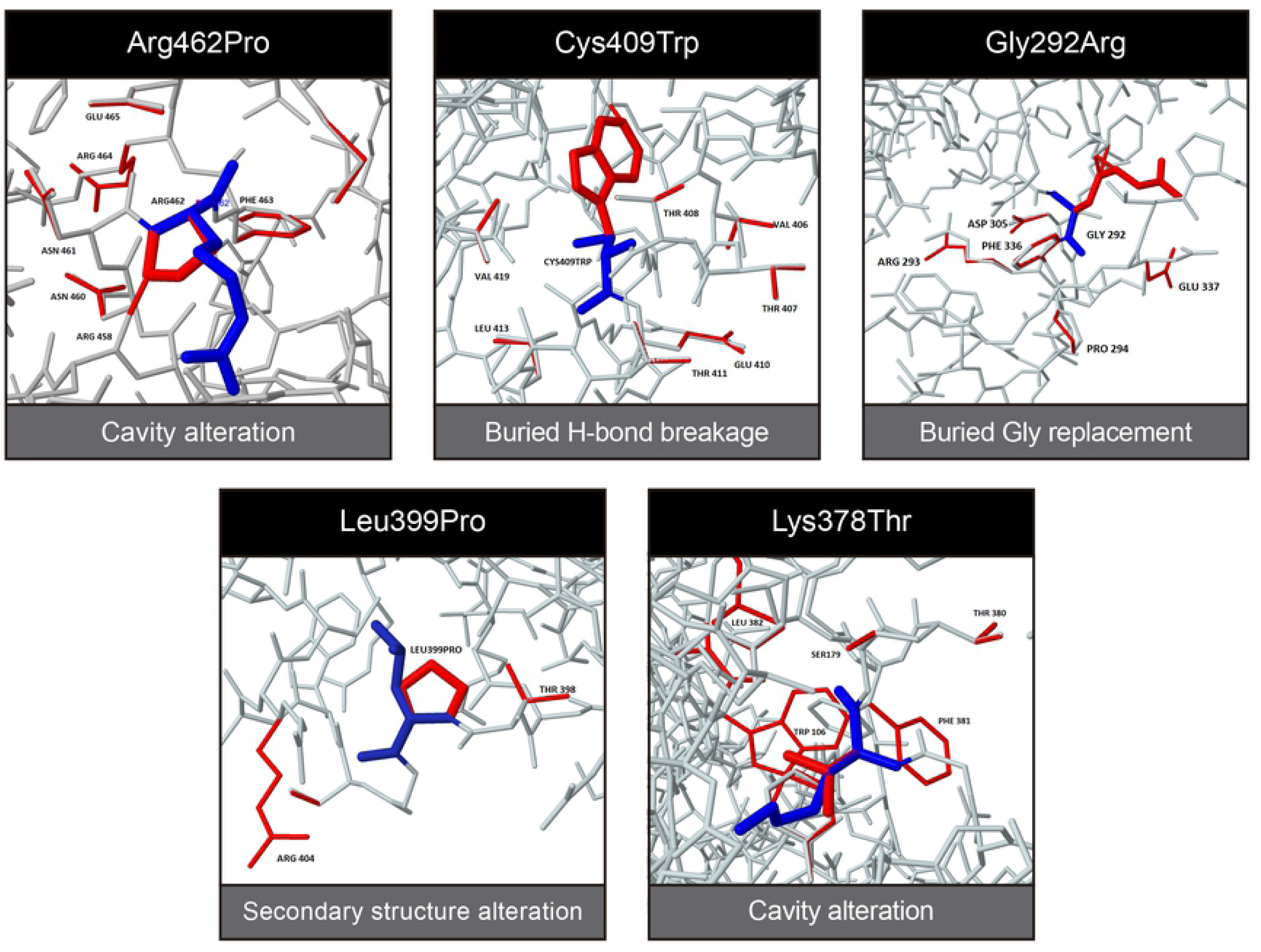
Missesnse3D structure prediction of changes generated by Single Amino acid Variants reported in patients with PNKP-associated diseases. In blue, Wild-Type residue is represented. In red, the variant generated by the mutation and the nearby residues affected in their space configuration.

#### Kinase domain

Although Arg462Pro is not located within a hotspot site, according to the Missense3D prediction, the variant generates an alteration that affects nearby amino acids (460,463,464) leading to the expansion of cavity volume by 117.72 Å^3. According to COACH prediction, these nearby amino acids are binding sites for ATP/ADP. The change in folding free energy (ΔΔG) is stabilizing. Lys378Thr is located within a hotspot associated with ATP/ADP binding site. Missense 3D predicted a contraction of cavity volume by 74.736 Å^3 and disruption of all side-chain/main-chain H-bond (s) formed by a buried Lys residue (RSA 3.4%). The protein is predicted destabilizing, increasing protein flexibility. Gly375Trp is also within a hotspot (ATP/ADP-binding site). However, it does not generate structural alteration according to Missense3D and the change in ΔΔG is predicted stabilizing, decreasing molecule stability (Table 1).

The rest of the mutations in the kinase domain do not present any clear tendency. Pro343Leu and Gly442Ser present low SuSPect score and intermediate conservation score. Neither generate structural alterations. Although the ΔΔG is predicted stabilizing for both Pro343Leu and Gly442Ser, the latter is associated with an increase in molecule flexibility. Leu399Pro changes ‘H’ (4-turn helix) to ‘T’ (hydrogen-bonded turn). Cys409Trp generates a structural alteration that disrupts all side-chain/main-chain H-bond (s) formed by a buried CYS residue (RSA 2.2 %) and the ΔΔG is stabilizing, decreasing molecule flexibility.

#### Phosphatase domain

Leu176Phe is located within a disease-propensity hotspot (DNA-binding site) but it does not generate structural damage and the ΔΔG is predicted to be stabilizing. Gly292Arg replaces a buried GLY residue (RSA 5.9%) with an exposed ARG residue (RSA 10.4%). Glu326Lys does not generate structural damage nor is it a ligand-binding site but the ΔΔG is predicted destabilizing, decreasing protein flexibility.

### PNKP mutations in gnomAD and genetic tolerance analysis

We also wondered if there were other mutations in PNKP not yet reported as clinical cases. We looked for PNKP mutations reported in the gnomAD v2.1.1 dataset which includes more than 141.000 individuals. We found a total of 424 missense mutations, but only 193 in the kinase domain, 132 in the phosphatase and 71 in the FHA domain (Fig. 11). Based on these results, observed mutations in the kinase domain are higher than expected compared to the phosphatase domain (d.f. = 1, □^2^ = 11.449; p < 0.001). Thus, the observed/expected ratio (O/E) for the kinase domain (1,18) is higher than the phosphatase domain (0,81). In addition, we found 12 *stops gain* mutations in the kinase and only 3 in the phosphatase. From the 31 frameshift variants, we observed 23 in the kinase and only 8 in the phosphatase. Of all missense variants, only 12 reported homozygous individuals, but they represented a very small portion since each variant has a high allelic count.

**Figure 11.**
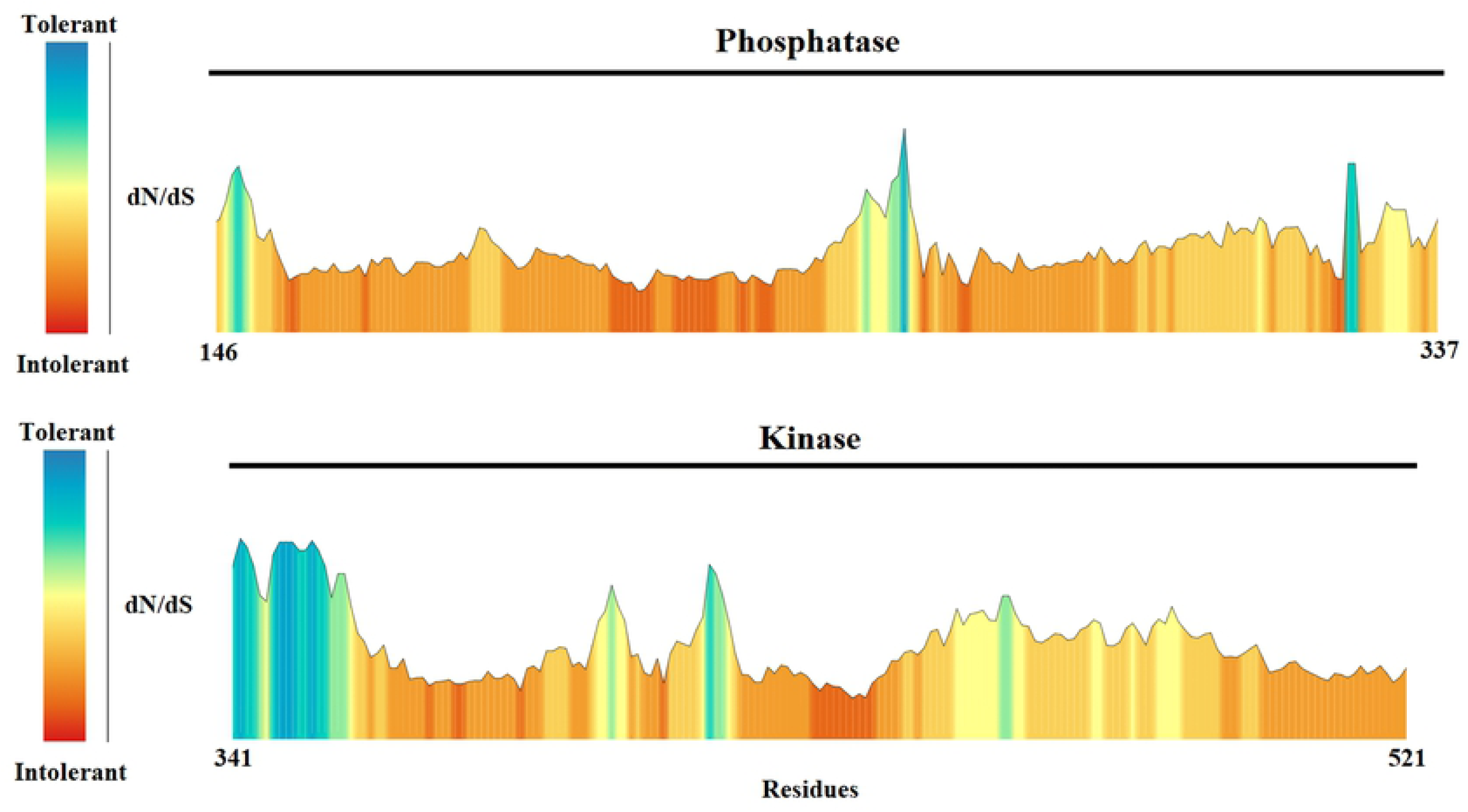
Mutational tolerance landscape of the phosphatase and kinase domain according to the dN/dS ratio. “Valleys” represent the most intolerant residues while “hills” represent neutral and tolerant residues.

Interestingly, there are two variants with a high number of homozygotes exclusive to Asia and others exclusive to Africa (supplementary figure 2). Of the 12 homozygous mutations found, 5 were in the phosphatase domain, but only one (Arg301Trp) is predicted to generate a structural alteration. For this variant, Missense3D predicted a cavity alteration, but this amino acid is not located within a hotspot. In addition, this mutation has a low SuSPect value (31) and the conservation score of amino acid 301 is very low (3). In general, the other mutations have low SuSPect values, as well as low conservation scores respectively: Tyr196Asn (16, 6), Arg139His (11, 3), Arg180Ser (18, 5), Glu508Lys (18, 5), Arg462Gln (13, 6), Ala441Gly (44, 6), Arg301Trp (31, 3), Thr217Ser (39, 9), Arg141Gln (11, 6), Gly244Arg (12, 4). Notably, most of the residues (except Thr217 and Ar462) are located in the surface of the protein (data not shown).

To obtain further information on the tolerability of PNKP mutations, we used the MetaDome server to compare the mutational tolerance landscape of each domain in terms of the nonsynonymous over synonymous ratio (dN/dS). We observed more valleys in the phosphatase domain related to more “intolerant” regions. In contrast, the kinase domain presented less “valleys” and more “hills”, relative to more “tolerant” regions. (Fig. 11). In fact, when we compared the ratios between domains, we found a higher tolerability in the kinase domain (W=21063, p-value = 0.0003958; supplementary figure 3). It was not surprising that when we analyzed both disease-propensity scores (SuSPect) and this tolerance landscape plot, there was an almost perfect overlap. For those intolerant dN/dS “valleys” the SuSPect score was the highest, compared to the “hills” that presented the lowest SuSPect score (Figure 12).

**Figure 12.**
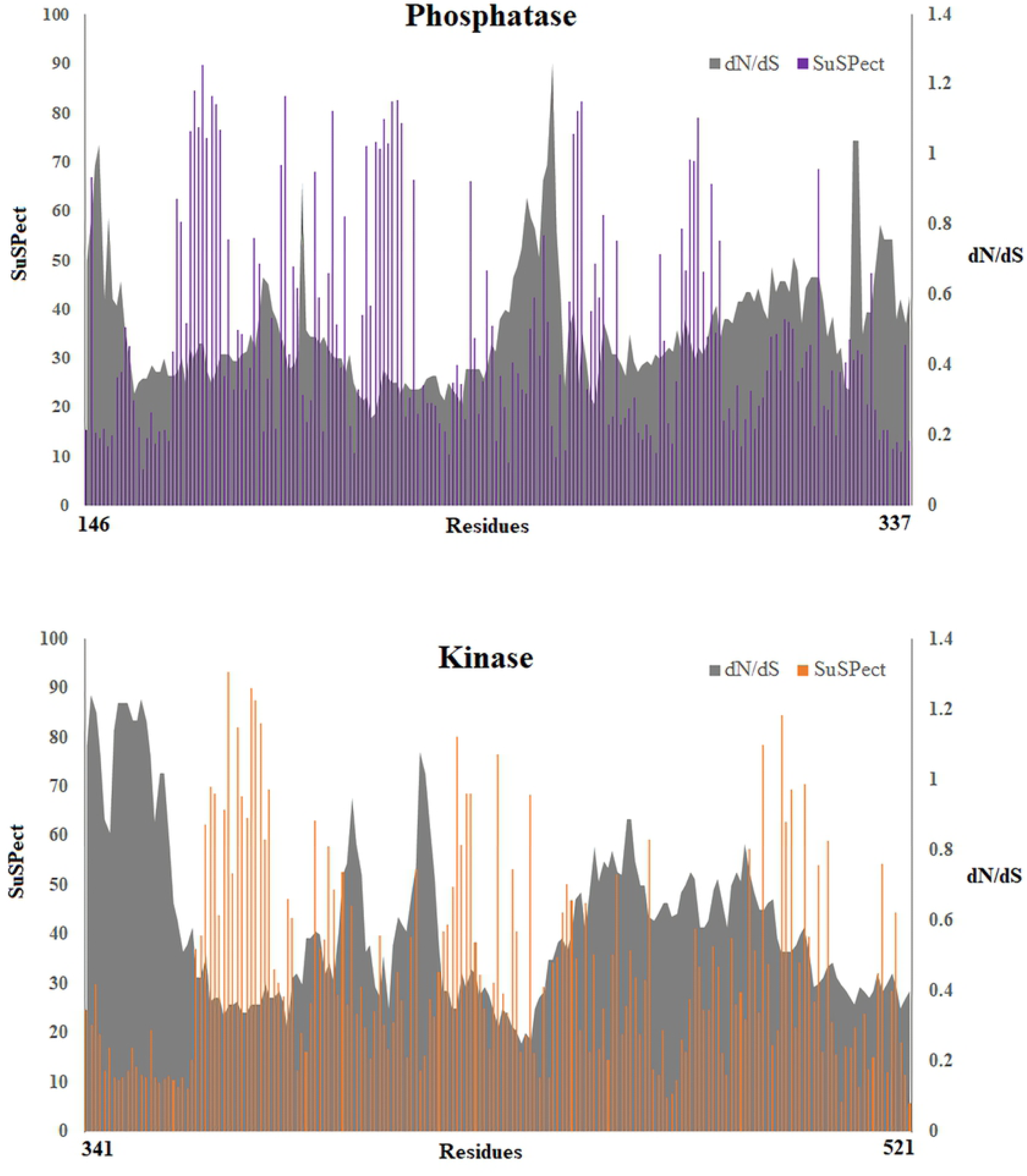
Mutational tolerance and disease susceptibility landscape in the phosphatase and kinase domain. The gray shaded areas represent the values of the dN/dS ratio and the corresponding mutational tolerability (the lower the value, the lower the tolerability). Bars represent SuSPect scores related to disease susceptibility (0-100). The “valleys” represent the residues with the lowest dN/dS ratio (intolerant), which show an overlap with the highest SuSPect scores.

## Discussion

### PNKP structural and evolutionary background

Current bioinformatics tools provide useful *in silico* analysis that combine theoretical-experimental knowledge to determine the relationship between protein structure, function, and disease. In our analysis, multiple sequence alignment was very informative to determine that human PNKP protein sequence has a low sequence identity among the analyzed species. As mammals represent more than 50% of the species analyzed, it would be expected that the percent identity would be very high in a DNA repair enzyme with a function as important as that of PNKP. Surprisingly, we found a large difference between the percent identity of PNKP with respect to POLB, PARP1, and LIGIII among species. Interestingly, like PNKP, the sequence of XRCC1 presented low identity, especially in non-mammals. This made us wonder if the degree of conservation of each domain of PNKP could be different. As we supposed, the phosphatase domain was more conserved as it presented more amino acids with higher conservation scores and less with low conservation scores. This could be related to the fact that the phosphatase sequence seems more conserved in different model organisms since orthologs are DNA phosphatases. In contrast, the kinase domain has enzymatic orthologs in different pathways. We believe that evolutionary rates in the kinase domain are therefore expected to be much higher.

As no crystal structure of the human PNKP has been deposited in the RCSB PDB database, we predicted the structure of PNKP with different servers in order to perform further structural analyzes. Although the Modeller software made the most accurate model, the structure assessment shows that it is still deficient in some rubrics, especially Ramachandran outliers. However, when we generated the structure of PARP1 with Modeller, we observed that the Ramachandran scores are comparable. This result is remarkable since structure of PARP1 has been completely characterized by different methods. In addition, the structural alignment showed that our model was comparable (very low RMSD) to the crystal phosphatase-kinase structure of the mouse.

### Prediction of pathogenicity of mutations in PNKP

The three mutations reported in the phosphatase domain are associated with the most severe phenotype (MCSZ), however, none are predicted to generate a significant structural alteration. As expected, Gly292Arg and Glu326Lys present low SuSPect values but the conservation scores are high. Gly292Arg has only been reported as a compound heterozygous together with Ala55Ser. This variant in the FHA domain has low SuSPect score (39) and is located in a conserved site (8) without prediction of structural damage. Leu176Phe mutation is within a disease-propensity hotspot and was reported to exhibit reduced levels of both DNA phosphatase and DNA kinase activity, at 30°C [30]. One may think that this mutation should not be viable, especially in a hotspot site, but it has only been reported in a heterozygous condition, together with Thr424GlyfsX48. As the latter variant only affects the kinase activity, we assume that it compensates for the affectation generated by Leu176Phe. In fact, it was proved that T424Gfs48X exhibited levels of DNA 3’-phosphatase activity comparable to WT despite having little DNA 5′-kinase activity [30]. Interestingly, the homozygous T424GfsX48 allele (causing MCSZ in humans) were lethal in mice at the embryonical level [31]. Thus, we believe that Leu176Phe mutation as others in PNKP can be under some degree of interallelic complementation as this is most often observed in multi-domain proteins [32]. However, Glu326Lys has only been reported in homozygous condition in severely affected individuals [20]. According to this, one may think that this mutation represents an exception for our proposal, but it is more a confirmation. It was demonstrated that E326K exhibited almost normal levels of both DNA 3′-phosphatase and DNA 5′-kinase activity at 30°C [30]. However, the mutation results in 10?to 20-fold reduced cellular levels of PNKP. In line with this, we found that this mutation is predicted to be destabilizing (ΔΔG = −0.788 kcal/mol). Thus, the severity of the phenotype generated by this mutation further confirms that insults in the phosphatase domain (especially in homozygous condition) are highly intolerant in terms of survival.

In the kinase domain, the Lys378Thr mutation affects the ligand-binding site and is predicted very pathogenic, only reported in MCSZ. Gly375Trp is located also in the ligand binding site (ATP/ADP) and although it does not generate structural alteration, it is predicted pathogenic as well. However, Gly375Trp has only been associated with AOA. In this case, as both can be in homozygous condition, the genotype, and the prediction of its effect on protein activity is not related to the observed phenotype. To support this, Arg462Pro also affects nearby residues of the ATP/ADP binding site in the kinase domain but can cause MCSZ when homozygous or AOA when accompanied by Leu399Pro. Thus, mutations in the active site of the kinase domain can not only be tolerated but can be found in a homozygous condition. Likewise, Leu399Pro, Cys409Trp and Pro343Leu mutations (all in the kinase domain) usually occur in a compound heterozygous condition, accompanied by a second allele with more serious insults in the kinase domain, usually deletions or frameshifts [22].

From all the mutations analyzed in gnomAD, the majority were located in the kinase domain. None of the homozygous mutations reported in the phosphatase domain are located in any hotspot. In addition, the general prediction for the homozygous mutations is that they are not pathological, possibly because most of them are located in the surface of the protein. This is very important since surface residues are less evolutionarily conserved and do not present a restricted conformational space, leading to potentially more viable physicochemical changes [7].

### Mutational Survivorship bias

Most of the disease-causing mutations in PNKP are confined to the kinase domain. However, not all combinations are equally pathogenic as many patients harboring only kinase mutations can present either severe (MCSZ) and/or milder phenotype (AOA/CMT). In contrast, only three mutations have been reported in the phosphatase domain and the three of them are highly pathogenic (MCSZ). A few novel mutations in the FHA domain are associated with both AOA and MCSZ [33,34]. Remarkably, most mutations in PNKP are presented as compound heterozygotes with only few mutations reported as recessive. This suggests that there is a gene dose effect on the pathogenicity of PNKP mutations. In fact, it has been determined that more deleterious mutations are more likely to be recessive than less deleterious mutations [35]. As dominance arises as a consequence of the functional importance of genes and their optimal expression levels, PNKP may be subject to interallelic complementation. In fact, E326K is the only recessive mutation in the phosphatase domain, but it has been shown to affect neither phosphatase nor kinase activity, but to decrease cellular levels of PNKP. The other mutations in phosphatase are accompanied by another mutation in the kinase that does not affect the phosphatase activity of the protein.

At the functional level, PNKP kinase and phosphatase activities do not act in concert on substrates containing both 5’-hydroxyl and 3’-phosphate termini, but PNKP phosphatase activity is much faster than the kinase activity. In addition, the 5’-phosphorylation of strand breaks containing a 3’-phosphate and 5’-hydroxyl depends on the preprocessing by the phosphatase activity of PNKP [36]. Likewise, it was demonstrated that the overexpression of wild-type recombinant PNKP, but not 3′-phosphatase-dead PNKP, can override the requirement for PNKP interaction with XRCC1 for rapid rates of SSBR following oxidative stress [37]. This can be related to the fact that PNKP preferentially binds to 3’-Phosphate substrates, so that the binding of the phosphatase domain to DNA blocks the additional DNA binding of the kinase domain [36]. The priority activity of the phosphatase domain over the kinase domain may lie in the relative importance of the two activities against the type of DNA damage, which is most often 3’-phosphate termini [38].

It has been shown that disease-resistant domains and proteins are more able to tolerate mutations, as they show significantly higher mutation rates and less degree of sequence conservation than disease-susceptible proteins and domains [3]. In line with this, the site-directed mutagenesis of the kinase domain active site does not affect the phosphatase activity of PNKP. In contrast, the disruption of the phosphatase domain also abrogates kinase function when a 3’-phosphate in the substrate is found [36]. It was suggested that this inhibition of the kinase domain is the result of steric hindrance by the phosphatase domain substrate rather than by allosteric restructuring of the enzyme. Finally, it has been shown that the presence of a 3’-Phosphate completely blocks the repair of DNA breaks because it is not processable by either DNA polymerase or ligase. In contrast, a 5’-hydroxyl only blocks the ligation and this is suggested to be bypassed (somehow) and therefore be more tolerated [36].

Enzymes from both the SSBR/DSBR pathway have mutations associated with pathological phenotypes like those associated with PNKP, but only PNKP mutations appear to combine both phenotypes (neurodevelopmental and neurodegeneration). The double function of PNKP in DNA Damage Response allows the enzyme to participate in both SSBR and DSBR, but for reasons that remain unknown, the elements that define which is the pathway that is most affected first at the intracellular level in the nervous system is what determines the appearance of the clinical manifestations. Like germline deletion of other key components of BER such as XRCC1, PNKP is essential for early embryogenesis [39]. Even when restricted to the nervous system, the deletion of PNKP still resulted in lethality [31]. Surprisingly, PNKP loss was substantially more severe than inactivation of either LIG4 (DSBR) or XRCC1 (SSBR) in mice, indicating the importance of this enzyme in repairing a broader range of DNA lesions than XRCC1 or LIG4 alone [31]. As neurons in the subventricular zone are reported to be very sensitive to the presence of unrepaired DSBs and readily undergo apoptosis, defective PNKP activity would lead to increased levels of apoptosis in the developing brain resulting in a reduced total brain volume as seen in MCSZ [40].

As none of the disease-causing mutations ablate DNA 3’-phosphatase activity of PNKP, it is possible that MCSZ cells could retain residual levels of PNKP [30]. This residual activity can be sufficient to keep the levels of DNA damage below the threshold for apoptosis in post-mitotic neurons, which are not as sensitive as newly differentiated neurons in the developing brain [40]. In fact, as previously suggested by our group [41], it was recently demonstrated that PNKP is important depending on cellular stage and tissue. In a recent work, Shimada and colleagues irradiated fibroblasts, induced-Pluripotent Stem Cells (iPSCs) and Neural Progenitor Cells (NPCs) at 5 Gy. One hour after irradiation, RNA samples were extracted and analyzed by RNA-Seq. Interestingly, iPSCs but especially fibroblasts showed little dependence on PNKP to repair DSBs via NHEJ [42]. Instead, the NPCs showed a high level of PNKP transcription both at baseline and after irradiation. This could explain recent findings suggesting that DSBR is not the cause of the neuropathology associated with PNKP-mutated diseases. In this experiment, authors only used patient-derived fibroblasts and were irradiated even at lower dose (2 Gy) [43]. Another experiment using PNKP-deficient HCT116 and HeLa cells generated with CRISPR/Cas9 showed that cells were biochemically competent in removing both protruding and recessed 3’-phosphates from synthetic DSB substrates [44]. However, the removal was much less efficiently than WT cells, so the authors suggested an alternative 3’-phosphatase. In a recent work, only MCSZ cell lines exhibit a defect in repair of IR-induced SSBs implicating reduced PNKP-dependent DNA phosphatase [43]. The authors concluded that it is reduced DNA 5’-kinase activity that is the major contributor and/or cause of the neurodegeneration in PNKP-mutated disease. This may be true if we assume that the kinase domain is the main contributor to the disease because it is the only one that can generate a pathological -but still viable-phenotype.

Thus, since the majority of mutations of the surviving patients are in the kinase domain, we propose that mutations in this domain represent “the areas where an airplane could suffer damage and still safely return” but the phosphatase domain, that is rarely mutated, represents the areas that when “attacked, would cause the plane to be lost” (Fig. 13). Even when all mutations are deleterious in the kinase domain, we find greater mutation tolerability in terms of survival. The kinase domain in general presents higher mutation rates in both clinical reports and gnomAD database as well as higher dN/dS ratios. It presents a lower degree of sequence conservation and fewer sites considered disease-propensity hotspots compared to the phosphatase domain. This means that the kinase domain can tolerate serious damage in places such as the active site and still generate a viable phenotype. However, very few patients are reported with damage in the phosphatase domain, as most mutations in this domain are very unlikely to be viable. This can be explained due the low mutation rates, higher degree of sequence conservation, lower dN/dS ratios, and more disease-propensity hotspots.

**Figure 13.**
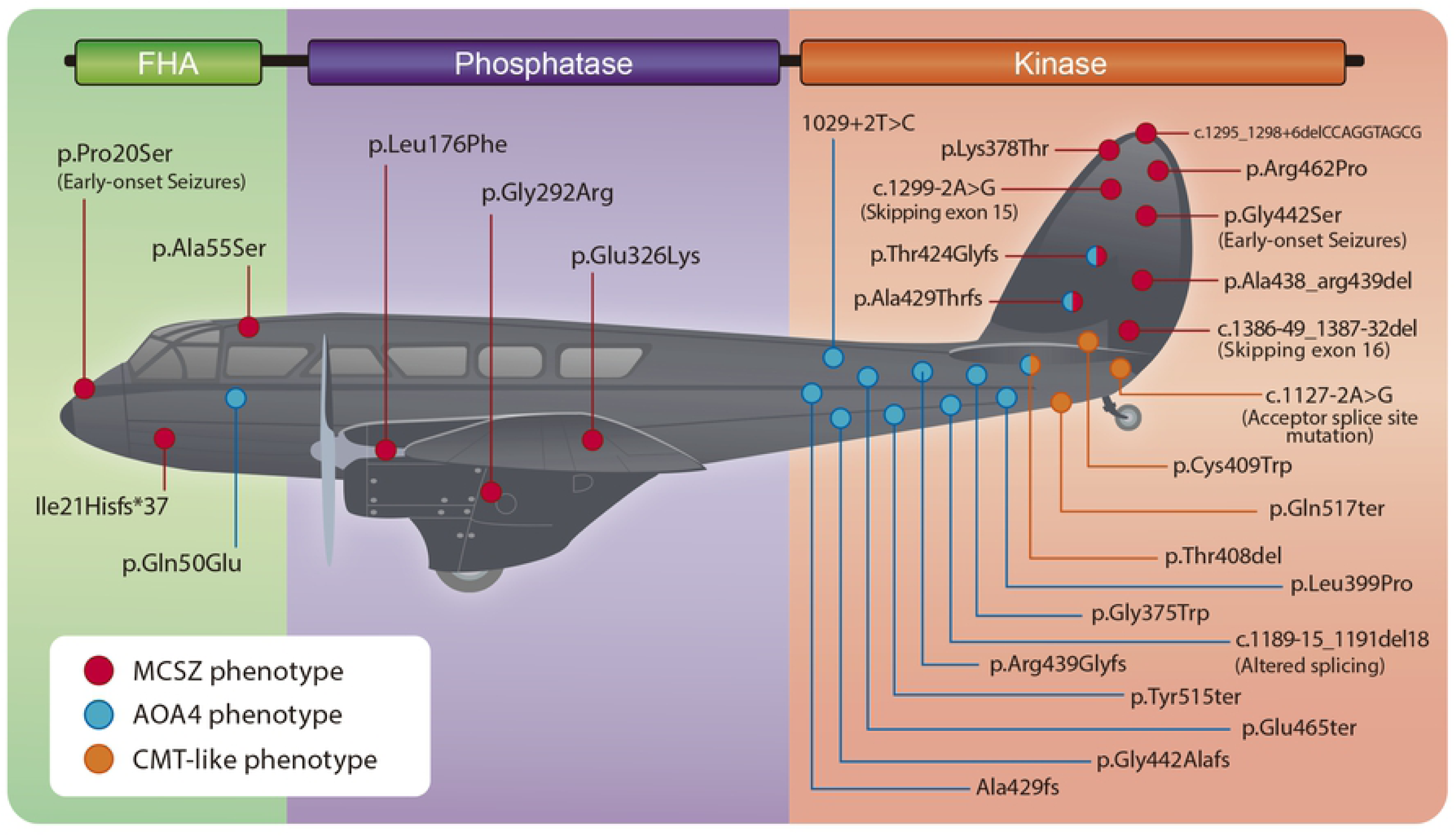
Depiction of PNKP domains as one of the airplanes from the Second World War from Wald’s analysis. The regions mutated within the kinase domain are the sites where the protein can be damaged and still generate a viable organism. In contrast, only three mutations have been reported in the phosphatase domain and patients have the worst phenotype: MCSZ. Therefore, the phosphatase domain represents the areas that when attacked, would cause the plane to be lost.

Finally, we would like to propose the term “Wald’s domain” for multi-domain proteins. For future studies, we do not propose to compare tolerability between proteins, because there are proteins that are intrinsically more important for the survival of the cell and/or organism. The idea is to compare the domains within one protein to identify whether one of them is under survivorship bias or not. In this case, mutations in the kinase are under survivorship bias, because it groups most mutations associated with disease, but is not the most critical for protein function and survival. Therefore, the domain with the lowest O/E mutation ratio, the highest degree of sequence conservation, the highest susceptibility to disease and the lowest dN/dS is a Wald’s domain. Thus, this approach constitutes a new way of evaluating the effect of mutations on the phenotype-genotype correlation in terms of survival.

## Conclusions

From Wald’s perspective, the regions mutated within the kinase domain are under survivorship bias since these sites represent points where the protein can be damaged and still generate a viable phenotype. It is also possible that the mutated domain is not related to a specific phenotype (MCSZ, AOA, CMT), although it could influence its pathogenicity. Together, our results support previous experimental evidence that demonstrates that the phosphatase domain is functionally more necessary and relevant for the repair of DNA lesions, especially in the context of the development of the central nervous system. Our data illustrate the value of available bioinformatics tools to determine the relationship between protein structure and function, as well as assessing whether a protein domain is under mutational survivorship bias or not. We suggest considering this bias when analyzing the role of different mutations in genotype-phenotype correlation.

## Methods

### Multiple sequence alignment and conservation analysis

Human PNKP protein sequence and other seventy sequences from different species were collected, sixty-three obtained from Uniprot (http://www.uniprot.org) [45] and seven obtained from NCBI Protein dataset (https://www.ncbi.nlm.nih.gov/protein). We chose the most representative species of each taxonomic group: 42 mammals, 8 bony fish, 5 reptiles, 3 amphibians, 3 birds, 2 arthropods, 2 apicomplexan, 1 amoebozoan, 1 cnidaria, 1 Mollusca, 1 bacterium and 1 virus (mimivirus). Uniprot: *Homo sapiens* (Q96T60), *Mus musculus* (Q9JLV6), *Rattus norvegicus* (A0A0G2JUH4), *Pan troglodytes* (K7BAU7), *Bos taurus* (F1N3Q9), *Anolis carolinensis* (G1KMT2), *Pongo abelii* (H2NZP9), *Ictidomys tridecemlineatus* (I3N446), *Otolemur garnettii* (H0WRT3), *Chlorocebus sabaeus* (A0A0D9S429), *Loxodonta africana* (G3TPU6), *Macaca fascicularis* (A0A2K5×601), *Saimiri boliviensis boliviensis* (A0A2K6TGX4), *Rhinopithecus bieti* (A0A2K6JZR0), *Colobus angolensis palliatus* (A0A2K5JFF3), *Propithecus coquereli* (A0A2K6FTG9), *Aotus nancymaae* (A0A2K5F2F6), *Cebus capucinus imitator* (A0A2K5R4×3), *Mesocricetus auratus* (A0A1U7R368), *Dipodomys ordii* (A0A1S3G4D4), *Mandrillus leucophaeus* (A0A2K5YE36), *Cavia porcellus* (H0V4R0), *Cricetulus griseus* (G3I715), *Myotis lucifugus* (G1PW59), *Ailuropoda melanoleuca* (G1M4Y9), *Capra hircus* (A0A452FU49), *Felis catus* (A0A337SAT2), *Sus scrofa* (F1RHU1), (*Ursus maritimus* (A0A452TNF4), *Ursus americanus* (A0A452QHV0), *Mustela putorius furo* (M3XT89), *Callithrix jacchus* (F6RVI4), *Drosophila melanogaster* (Q9VHS0), *Danio rerio* (Q08BP0), *Oryzias latipes* (H2MLN4), *Xenopus tropicalis* (B1WBK3), *Equus caballus* (F6QG90), *Xenopus laevis* (A9UMQ0), *Canis lupus familiaris* (E2R0U3), *Fopius arisanus* (A0A0C9R1P0), *Latimeria chalumnae* (H3B245), *Electrophorus electricus* (A0A4W4HJQ6), *Acanthamoeba castellanii* str. (L8GVC5), *Nothobranchius furzeri* (A0A1A7ZB78), *Alligator sinensis* (A0A1U7R7V0), *Austrofundulus limnaeus* (A0A2I4BRQ3), *Balaenoptera acutorostrata scammoni* (A0A384ACF1), *Ictalurus punctatus* (A0A2D0SWM3), *Gorilla gorilla gorilla* (G3R3W6), *Acanthamoeba polyphaga mimivirus* (Q5UQD2), *Nothobranchius kuhntae* (A0A1A8JP17), *Physeter macrocephalus* (A0A455AU26), *Alligator mississippiensis* (A0A151NJZ6), *Erinaceus europaeus* (A0A1S3AK18), *Hydra vulgaris* (T2M7D9), *Castor canadensis* (A0A250XY04), *Plasmodium malariae* (A0A1A8W4×2), *Neovison vison* (U6D1K9), *Callorhinus ursinus* (A0A3Q7NQ21), *Plasmodium gallinaceum* (A0A1J1GNL7), *Mycobacterium* sp. (A0A1A0×7E1), *Odobenus rosmarus divergens* (A0A2U3WM26), *Tarsius syrichta* (A0A1U7TJR7), *Vulpes vulpes* (A0A3Q7U9N3), *Delphinapterus leucas* (A0A2Y9P3J8). NCBI: *Gopherus evgoodei* (XP_030400898.1), *Microcaecilia unicolor* (XP_030053814.1), *Podarcis muralis* (XP_028557931.1), *Falco cherrug* (XP_027653441.1), *Lonchura striata domestica* (XP_021405459.1), *Gallus gallus* (XP_025001234.1), *Aplysia californica* (XP_005090552.1).

Multiple alignment of these sequences was performed with EMBL-EBI’s Multiple Alignment using Fast Fourier Transform (MAFFT) platform with default settings [46]. We performed a test using the BlastP tool [26] to obtain the percent identity between the human PNKP sequence with respect to the rest of the species used in the alignment.

### Percent identity analysis of SSBR proteins

The human protein sequences of PNKP, PARP1, XRCC1, PARP1, and LIG3 were obtained in Uniprot [45] (Table 2) and we performed a test using the BlastP tool [26] to obtain the percent identity of each human sequence with respect to seven different species (*Pan troglodytes, Equus caballus, Sus scrofa, Canis lupus, Felis catus, Mus musculus, Rattus norvegicus, Gallus gallus, Xenopus laevis, Danio rerio, Drosophila melanogaster*). The refseq_protein database (Max target sequences = 1000) was used and the sequences with the lowest E value, highest coverage and the highest similarity were chosen for each species.

**Table 2.** Sequence ID used for the analysis of percent identity of each SSBR protein.

| Species | POLB | PNKP | XRCC1 | LIGIII | PARP1 |
| --- | --- | --- | --- | --- | --- |
| *Pan troglodytes* | XP_009453<br>547.1 | XP_016792<br>104.1 | XP_016791<br>569.1 | XP_009430<br>685.3 | XP_003949<br>754.2 |
| *Equus caballus* | XP_001499<br>852.3 | XP_001917<br>391.2 | XP_001499<br>917.3 | XP_023508<br>784.1 | XP_023488<br>446.1 |
| *Sus scrofa* | XP_020933<br>913.1 | XP_005664<br>806.3 | XP_013844<br>100.1 | XP_020921<br>813.1 | XP_003357<br>689.2 |
| *Canis lupus* | XP_005629<br>859.1 | XP_005616<br>345.1 | NP_001273<br>911.1 | XP_005624<br>869.1 | XP_022277<br>419.1 |
| *Felis catus* | XP_003984<br>799.1 | XP_023101<br>490.1 | XP_003997<br>757.1 | XP_003996<br>610.2 | XP_003999<br>138.3 |
| *Mus musculus* | NP_035260.<br>1 | XP_006541<br>101.2 | NP_033558.<br>3 | NP_001278<br>175.1 | NP_031441.<br>2 |
| *Rattus norvegicus* | NP_058837.<br>2 | NP_001004<br>259.1 | XP_017445<br>286.1 | XP_008766 | NP_037195.<br>1 |
|  |  |  |  | 242.1 |  |
| *Gallus gallus* | XP_024998<br>666.1 | XP_025001<br>234.1 | XP_024997<br>899.1 | NP_001006<br>215.2 | NP_990594.<br>2 |
| *Xenopus laevis* | NP_001081<br>643.1 | NP_001108<br>303.1 | NP_001080<br>711.1 | NP_001082<br>183.1 | NP_001081<br>571.1 |
| *Danio rerio* | NP_001003<br>879.2 | NP_001071<br>046.1 | NP_001003<br>988.2 | NP_001025<br>345.2 | NP_001038<br>407.1 |

The difference in the sequence percent identity was calculated and an ANOVA test was used to determine whether there were differences between the proteins, while also considering the species as a possible source of variability that should be accounted for.

### Conservation analysis

The MAFFT method was used as it is recommended for better accuracy of multiple sequence alignments [47]. This alignment was used as input to estimate and visualize the evolutionary conservation of the amino acid sequence of human PNKP on the ConSurf 2016 platform [48]. The analysis performed with default settings and PNKP’s 3D structure prediction was generated using Modeller within the ConSurf server. The conservation scores for the amino acids of the phosphatase (146-337) and kinase (341-521) domain were analyzed and a homogeneity chi-squared test was performed to determine if the conservation values were different or not in each domain.

### PNKP 3D structure prediction

As there is no complete 3D structure of PNKP protein in the RCSB Protein Data Bank (only the FHA domain based on the mouse’s structure) different software was used to predict its 3D structure. We used Modeller [49], Swiss-Model [50], Phyre2 [51], and I-TASSER [52]. The modeling quality comparison was performed using the platform MolProbity [53], Structure Assessment tool of Swiss-Model. Final molecular graphics were performed with UCSF Chimera, developed by the Resource for Biocomputing, Visualization, and Informatics at the University of California, San Francisco [54]. Structure alignment was performed with TM-align server [55] using the structure from Modeller and the crystal structure of murine phosphatase-kinase domain (PDB ID: 3ZVL).

### Prediction of pathogenicity of mutations in PNKP

Based on the structure generated in Modeller, we use the SuSPect platform[18] to analyze the mutational sensitivity of the phosphatase and kinase domain amino acids. This method (Disease-Susceptibility-based SAV Phenotype Prediction) predicts how likely single amino acid variants (SAVs), are to be associated with the disease. We also analyze the conservation score of amino acids at the sites of the sequences where PNKP mutations of the data obtained with ConSurf occur. In addition, the Missence3D program was used [56] to predict the structural changes introduced by an amino acid substitution based on the most common PNKP mutations.

For the analysis of disease-propensity hotspots in the sequence of PNKP, the probability for a mutation in each position was calculated, for a total of 20 probabilities for each position. Furthermore, for determining said hotspots, the average of the 20 probabilities was calculated for each amino acid, and then compared to a predetermined value, in this case, the 95th percentile, to establish the positions that had a higher average mutation susceptibility compared to the rest of positions. Finally, to eliminate possible single positions with abnormal probabilities, only consecutive positions were used to determine the existence of these disease-propensity hotspots.

### PNKP ligand-binding sites prediction

To confirm if disease-propensity hotspots correspond to functional ligand binding sites in PNKP we used COACH [27] and COFACTOR [28]. In addition, the Missence3D program was used [56] to predict the structural changes introduced by an amino acid substitution based on the most common PNKP single-nucleotide variation reported in patients. We analyze the potential post-translational modification (PTM) sites of PNKP by using NetPhos 3.1 [57]. To confirm these analyzes, we reviewed the databases Phospho.ELM [58] and PhosphoSitePlus v6.5.9.1 [59] that contain in vivo and in vitro phosphorylation data.

### Structural damage of PNKP mutations

We evaluated the structural damage of missense mutations reported in PNKP on the Missense3D platform [56] using the 3D structure predicted with Modeller. Dynamut [60] were also used to analyzed the impact of mutations on protein dynamics and stability resulting from vibrational entropy changes.

### PNKP mutations in gnomAD and genetic tolerance analysis

All reported (non-intronic) mutations for PNKP collected from both exome sequencing and whole genomes in the gnomAD v2.1.1 dataset [61] (https://gnomad.broadinstitute.org/) were analyzed. We only included variants from the canonical transcript (ENST00000322344.3). We especially analyzed “Missense” mutations to compare the incidence between domains and analyze those cases reported as homozygous. Low confidence variants were not considered. To obtain further information on genetic tolerance of mutations in PNKP, we used the MetaDome server [62] to plot the mutational tolerance landscape of each domain. Mann–Whitney U test will be used to compared the average dN/dS ratio between domains.

### Homology analysis of the phosphatase and kinase domains

The BlastP tool was used to analyze only the phosphatase (146-337) and kinase (341-521) domain sequence against the Protein Data Bank (PDB) and Uniprot/Swissprot database. A phylogenetic tree was created with the Blast Tree View tool.

## Funding

Support from the University of Costa Rica is gratefully acknowledged (project 111-B8– 372)

## Author contributions

LBG: conception and design of the work, analysis, and interpretation of data. Wrote the manuscript. GJH: Analysis, figures design, interpretation of data. AA: Statistical analysis, interpretation of data and manuscript revision. AL: managed the project, interpretation of data and manuscript revision.

## Declaration of interests

Authors declare no conflict of interest.

## Data Accessibility

The data that support the findings of this study are openly available in Figshare at 10.6084/m9.figshare.12644000 (not public yet).

For review purposes: https://figshare.com/s/ea69e39896793707f237

## Supplementary information

Supplementary Figure 1. BlastP phylogenetic tree of the sequence of each domain. If the BlastP analysis if performed with only the sequence of the phosphatase domain (146-337) against the “Swissprot” database, most significant alignments are with polynucleotide 3’-phosphatase proteins from different kingdoms: plants, fungi and animals. When performing the same analysis with the kinase domain sequence (341-521) against the “Swissprot” database, the greatest number of results are for Intron maturase (Maturase K). However, if the sequences from the Protein Data Bank (PDB) are used, at least 5 different proteins show a certain degree of homology with the sequence, mainly adenylate kinase.

Supplementary Figure 2. Number of Missense mutations in PNKP per domain reported in the gnomAD v2.1.1 dataset. Twelve mutations were found in homozygous condition in specific regions in the world.

Supplementary Figure 3. Comparison of the dN/dS ratio between the phosphatase and kinase residues. The average ratio for the kinase is higher, meaning this is a more tolerant domain (W = 21063, p-value = 0.0003958).

Supplementary Table 1. Evaluation of each 3D model using

